# Peak Space Motion Artifact Cancellation Applied to Textile Electrode Waist Electrocardiograms Recorded During Outdoors Walking and Jogging

**DOI:** 10.1101/2022.01.07.475456

**Authors:** Bruce Hopenfeld

## Abstract

**Background:** Obtaining reliable rate heart estimates from waist based electrocardiograms (ECGs) poses a very challenging problem due to the presence of extreme motion artifacts. The literature reveals few, if any, attempts to apply motion artifact cancellation methods to waist based ECGs. This paper describes a new methodology for ameliorating the effects of motion artifacts in ECGs by specifically targeting ECG peaks for elimination that are determined to be correlated with accelerometer peaks. This peak space cancellation is applied to real world waist based ECGs.

**Algorithm Summary:** The methodology includes successive applications of a previously described pattern-based heart beat detection scheme (Temporal Pattern Search, or “TEPS”). In the first application, TEPS is applied to accelerometer signals recorded contemporaneously with ECG signals to identify high-quality accelerometer peak sequences (SA) indicative of quasi-periodic motion likely to impair identification of peaks in a corresponding ECG signal. The process then performs ECG peak detection and locates the closest in time ECG peak to each peak in an SA. The differences in time between ECG and SA peaks are clustered. If the number of elements in a cluster of peaks in an SA exceeds a threshold, the ECG peaks in that cluster are removed from further processing. After this peak removal process, further QRS detection proceeds according to TEPS.

**Experiment:** The above procedure was applied to data from real world experiments involving four sessions of walking and jogging on a dirt road for approximately 20-25 minutes. A compression shirt with textile electrodes served as the ground truth recording. A textile electrode based chest strap was worn around the waist to generate a single channel signal upon which to test peak space cancellation/TEPS.

**Results:** Both walking and jogging heart rates were generally well tracked. In the four recordings, the percentage of segments within 10 beats/minute of reference was 96%, 99%, 92% and 96%. The percentage of segments within 5 beats/minute of reference was 86%, 90%, 82% and 78%. There was very good agreement between the RR intervals associated with the reference and waist recordings. For acceptable quality segments, the root mean square sum of successive RR interval differences (RMSSD) was calculated for both the reference and waist recordings. Next, the difference between waist and reference RMSSDs was calculated (ΔRMSSD). The mean ΔRMSSD (over acceptable segments) was 4.6 m, 5.2 ms, 5.2 ms and 6.6 ms for the four recordings. Given that only one waist ECG channel was available, and that the strap used for the waist recording was not tailored for that purpose, the proposed methodology shows promise for waist based sinus rhythm QRS detection.

## 1. Introduction

Motion artifacts remain a major problem for the detection of heartbeats in ambulatory electrocardiograms, especially with textile electrode based recordings[1, 2]. The difficulty is particularly severe in recording from the waist, a desirable location from the standpoint of comfort and convenience [3, 4].

Prior methods for reducing motion artifact include denoising an ECG signal with an adaptive filter [1] and through elimination of wavelet coefficients[5]. In either case, the goal of these techniques is to recover a denoised version of the ECG signal.

The recovery of a reasonably denoised signal is difficult if not impossible in the presence of a signal, such as that shown in the upper right hand panel in Figure 1, that is contaminated with high amplitude motion artifacts that can occur very close in time to true QRS complexes. Indeed, in such cases, RR interval/rhythm information may be the only information that can possibly be obtained from the ECG signal. Instead of attempting to recover some version of the original signal, the present paper describes a method for ameliorating the effects of motion artifact peaks so that the RR interval/rhythm information can be derived from the ECG signal.

**Figure 1.**
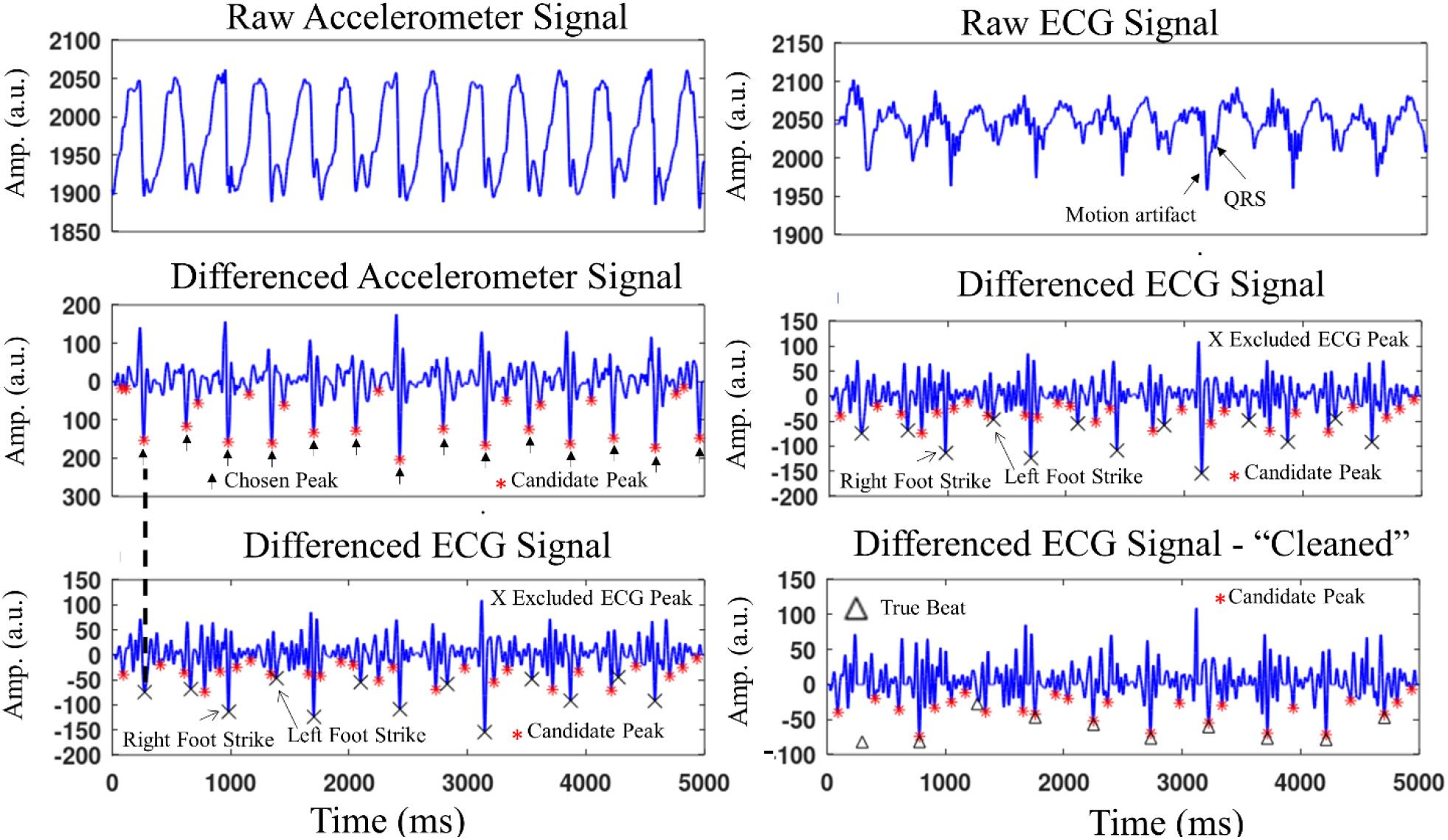
Simultaneous accelerometer and ECG signals. A high quality sequence is detected in the accelerometer signal (middle left panel) and correlated with peaks in the differenced ECG signal (lower left panel). These peaks are excluded from further ECG processing, as indicted by the X’s, resulting in (effectively) the ECG signal in the lower right panel. The differenced signal in the lower left panel is the same as that in the middle right panel.

Hopenfeld describes a temporal pattern searching methodology, TEPS, that takes advantage of the relative regularity of sinus rhythm to detect QRS complexes in noisy ECG signals[6, 7]. If a motion (e.g. jogging) results in a motion artifact sequence that is temporally regular, TEPS may produce a false positive detection of the artifact sequence. Elimination of such sequences is the main goal of the methodology described in this work.

## 2. Algorithm

Figure 2 is a high-level block diagram of the motion artifact cancellation scheme, which processes data in non-overlapping five second segments. For each segment, each of the three (x, y, z axis) accelerometer signals is first processed according to the TEPS methodology. Specifically, after pre-processing, the algorithm selects a certain number of peaks (“Candidate Peaks”) in each accelerometer signal, and generates a set of high quality sequences (SA) therefrom, where quality indicates the likelihood that a particular sequence corresponds to a repetitive motion (e.g. foot strikes while walking). The peak times of these sequences, as well as their average interpeak interval (1/cadence), are stored.

**Figure 2.**
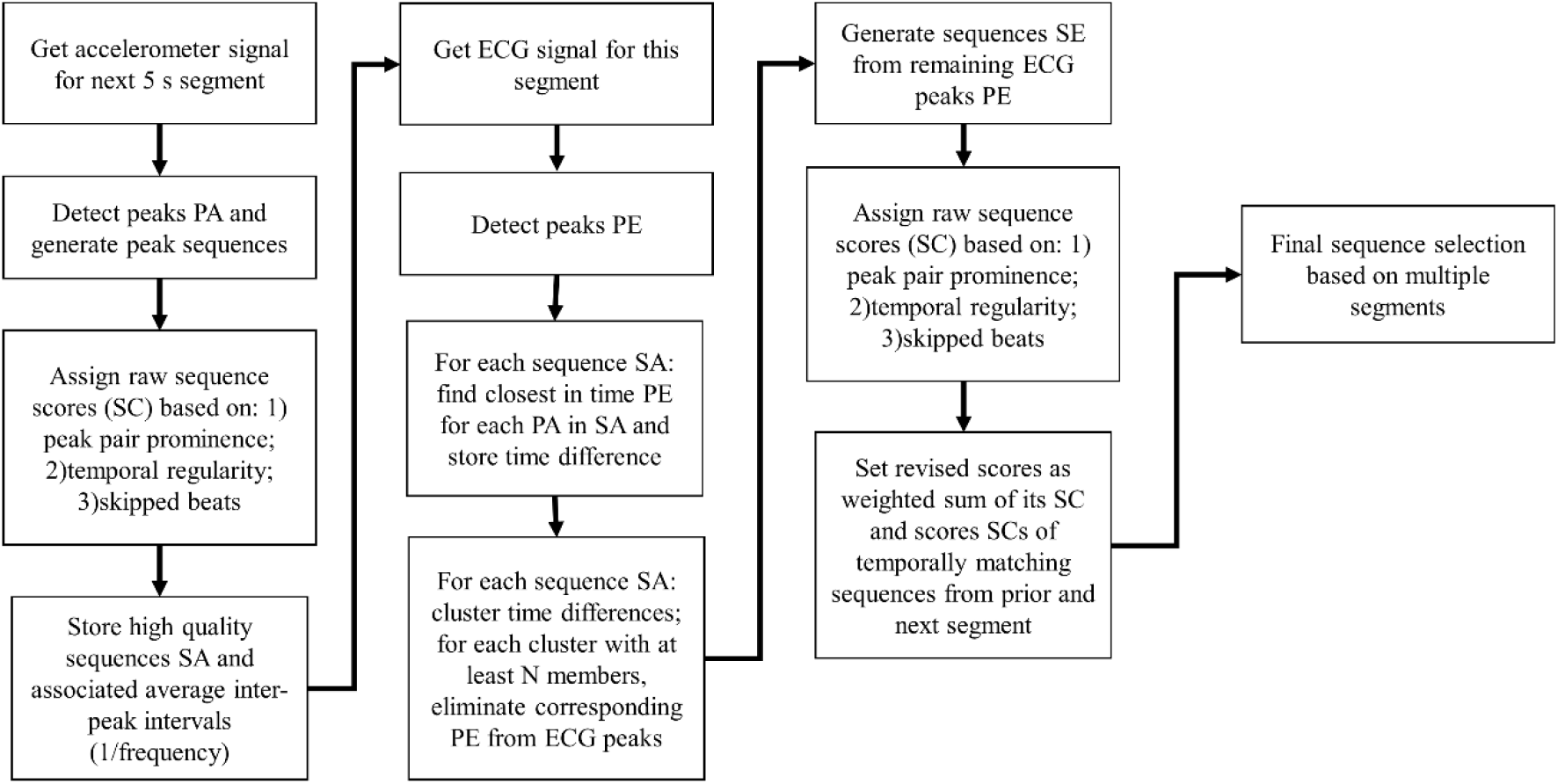
High level flowchart of peak space exclusion and single channel TEPS.

**Figure 3.**
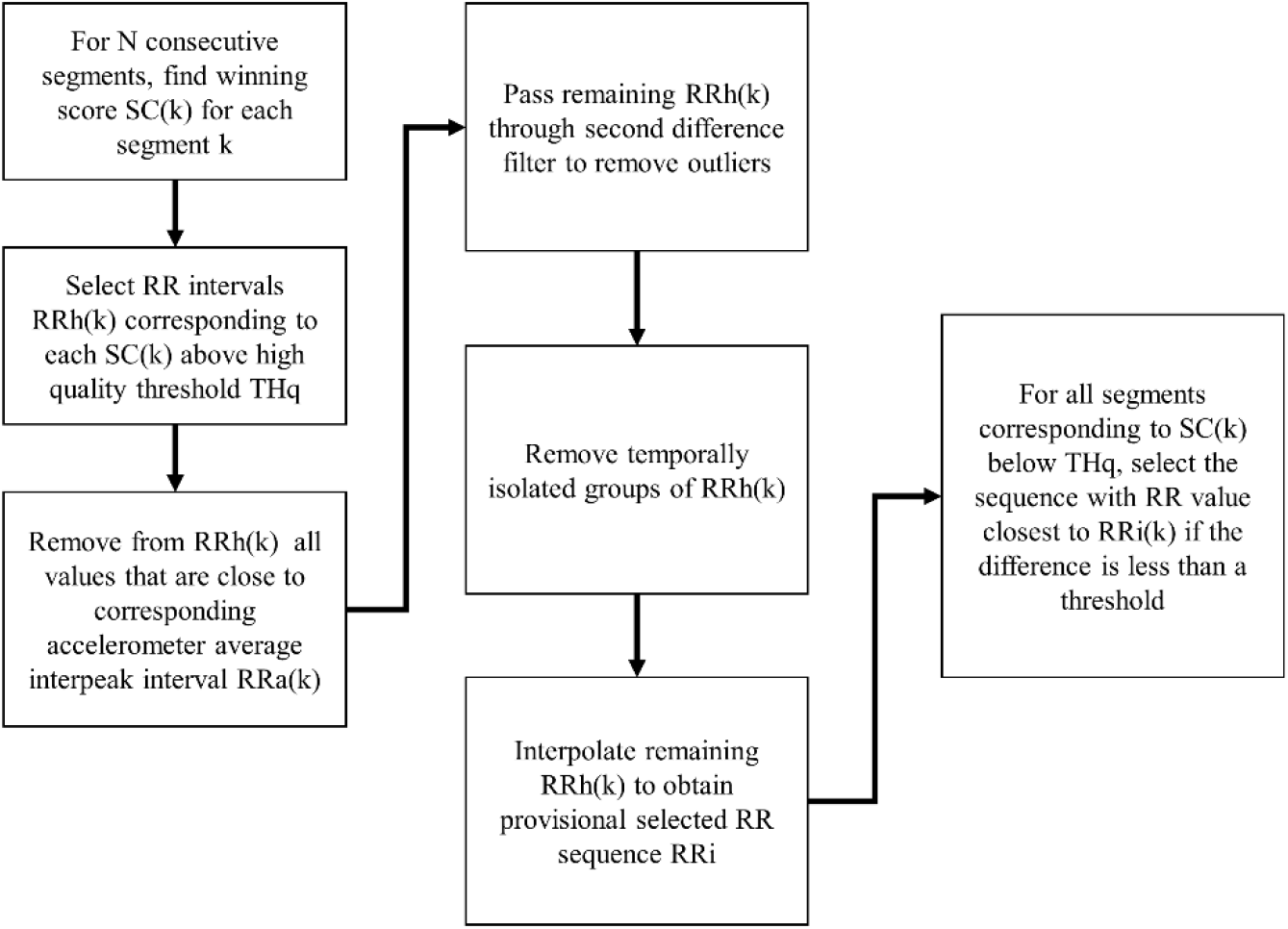
Flowchart of the multi-segment final sequence/RR value selection.

Next, the ECG signals are processed. An initial set of peaks is detected. For each of the accelerometer sequences SA, the closest in time ECG peak is located, thereby generating a set of peak time differences. The set of peak time differences is (one-dimensionally) clustered. Any cluster that has at least a prespecified number of entries (e.g. 3), is considered to be correlated to the accelerometer sequence, and the ECG peaks in that cluster are excluded from further processing.

ECG sequences are generated and scored as previously described. (If multiple ECG channels are available, the sequence scores are further based on peak timing coherence as described in [6].) This processing results, for each segment, in a set of sequences and associated average RR intervals and scores. The final selection of the best sequence for each segment depends on a multi-segment process that is based upon sequence quality scores and intersegment RR smoothness, while taking into account cadence information. Thus, the accelerometer signals are used for both motion artifact elimination, and for a higher level RR interval trajectory analysis. (This higher level analysis is necessary because the motion artifact elimination does not ensure that all motion artifacts are removed, which means that artifact based ECG sequences may still be generated.)

According to TEPS, both accelerometer and ECG sequences are scored for quality according to: 1) prominence of the peaks within a sequence; 2) temporal regularity; and 3) the number of skipped beats. If more than one ECG channel is available, ECG sequences are further scored according to the timing coherence of peaks between channels. The temporal regularity score component is a measure that quantifies the change in between peak time intervals over a sequence. (The time between a sequence’s consecutive peaks will be referred to as an “RR interval.”)

The raw sequence score SC is set equal to:

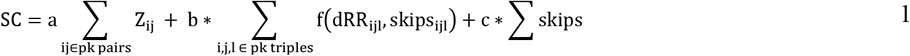

where Z is the prominence ratio for each peak pair in a sequence, dRR_ijl_ is the change in RR intervals between 3 consecutive peaks with corresponding skipped beats skips_ijl_, f() is a Gaussian with a standard deviation that depends on skips_ijl_. Additional details are described in [6]. The coefficients a, b and c may be chosen by adding calibrated amounts of noise to a signal with known peak locations and performing a least squares fit M * [a b c]’ = Q, where M is a matrix with the three scores described above and Q is a quality measure (e.g. F1) of the sequence corresponding to a row on the left hand side of the above equation. A more elaborate fitting procedure involving neural networks is also possible.

### 2.1. Accelerometer Peak Sequence Generation

The upper left panel in Figure 1 shows a raw accelerometer signal, upsampled from 100Hz (original recording) to 256Hz after low pass filtering with a 5^th^ order Butterworth filter with a cutoff frequency of 45Hz. The filtered signals is differenced according to: y(i) = x(i-12)-2*x(i-6)+x(i), which results in the signal in the middle left panel. The N (=30) peaks with the greatest prominence are chosen as Candidate Peaks, shown in the middle left panel in Figure 1, from which sequences are generated. Sequences are generated by generating all possible subsequence combinations in each of the two subsegments, eliminating sequences that do not meet physiological interpeak interval and temporal regularity criteria.

Sequences are scored according to Equation 1. The temporal regularity coefficient b was reduced by a factor of ¾ compared to ECG sequence scoring to allow for the greater irregularity of running/walking cadences compared to heart rate. Scores above a heuristically chosen threshold are tagged as high quality sequences, which served as the basis for ECG peak removal as will be further described below. Arrows in the middle left panel of Figure 1 point to a high quality sequence.

### 2.2. ECG Peak Sequence Generation/Artifact Sequence Exclusion

The raw ECG signal (Figure 1, right upper panel) is upsampled from 250Hz to 256Hz after filtering identical to the accelerometer raw signal. The filtered signal is similarly differenced (Figure 1, right middle panel, which shows the same signal as in Figure 1, left lower panel). For each high quality accelerometer sequence, the time delay between each peak in that sequence and the closest ECG peak is computed, yielding a set of time delay values DT_s_, where the s subscript indicates a particular accelerometer sequence. The values in DT_s_ are clustered, and if the number of elements in any cluster exceeds a threshold (e.g. 3), then the ECG peaks corresponding to the cluster are removed.

For example, for every peak in the high quality sequence shown in the middle panel of Figure 1, the closest ECG peak in the lower left panel of Figure 1 is located. In this example, every other high quality accelerometer peak, starting with the first one indicted by the dashed line connecting the middle and lower panels in Figure 1, had a corresponding ECG peak that occurred within −5 ms and 5 ms of the accelerometer peak. (Only 1 of the corresponding ECG peaks occurred before the accelerometer peak; the ECG peaks will generally occur after the accelerometer peak.) There were six ECG peaks in this cluster, as shown in Figure 4, where this cluster is indicated by the label “Right Foot Strike Cluster.” The other accelerometer peaks were associated with a 6 element cluster that had a delay between 20 ms, and 40 ms, cluster is indicated by the label “Left Foot Strike Cluster.” (Two peaks within this delay period were not included in the cluster because their prominences failed to meet a threshold; these peaks were too “small” to be Candidate Peaks anyway.) All 12 ECG peaks in the two clusters are excluded from the set of ECG candidate peaks, and further excluded from all further peak prominence calculations.

**Figure 4.**
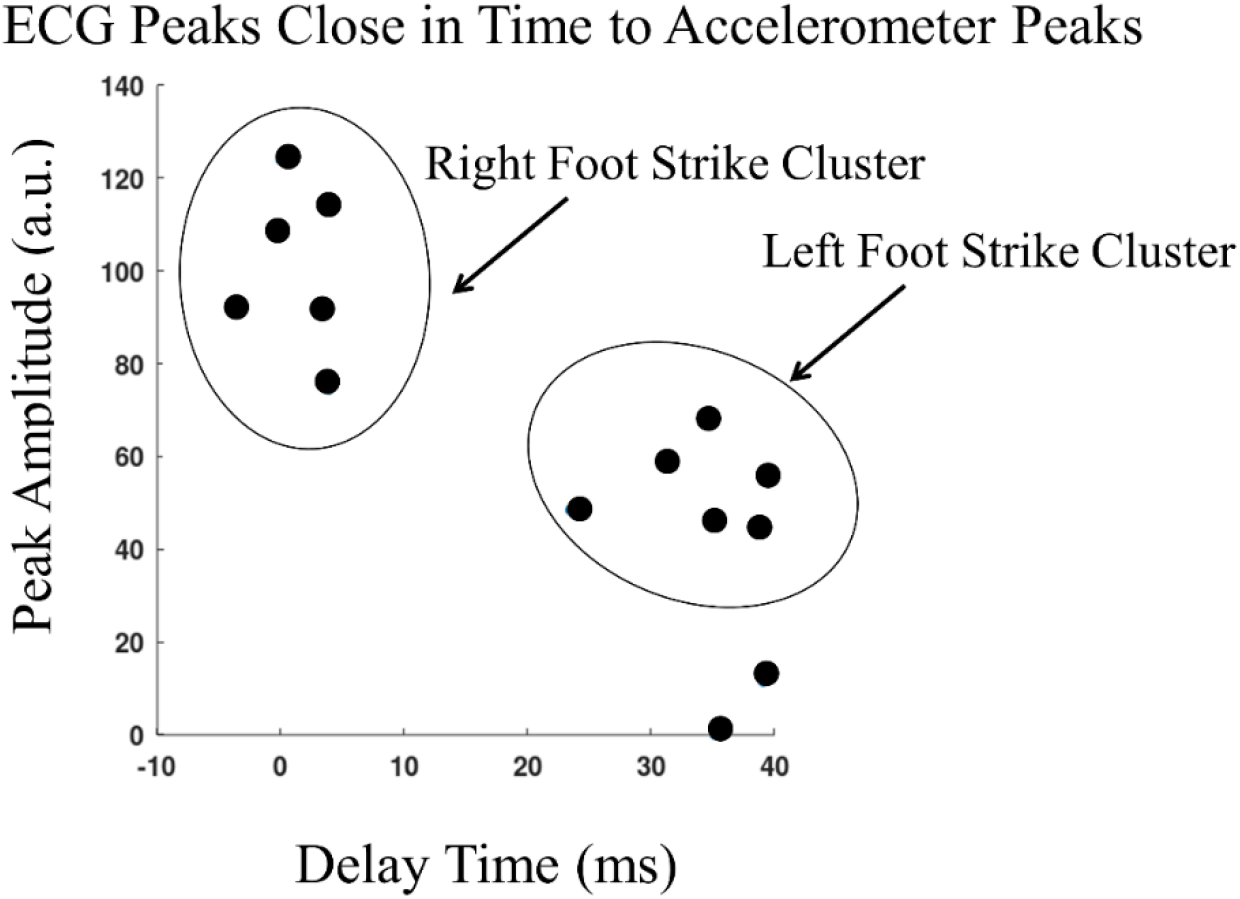
Clustering of the accelerometer related ECG peaks shown in Figure 1. In the present work, clustering is done solely in the time domain. This figure suggests the possibility of two dimensional clustering in future versions.

Thus, in effect, after the exclusion, the algorithm processes a signal shown in the bottom right panel in Figure 1. According to TEPS, each ECG sequence is scored by a peak pair prominence measure, which is the ratio of the amplitudes of the two sequence peaks to the amplitudes of intervening peaks. By eliminating the large amplitude artifact peaks, the peak pair prominence score for all ECG sequences is enhanced, which in turn provides greater confidence that a particular ECG sequence reflects true heart beats.

### 2.3. Final Sequence Selection/Multi-Segment Processing

Final sequence selection is based upon processing segments after they are acquired. However the process can be modified for real time sequence selection. Indeed, during real time processing, the detection of a high quality segment could constrain the search in the following segment and thus increase the efficiency of the sequence generation process.

The winning score SC(k) for each segment k is compared to a quality threshold THq. For those segments k with SC(k) > THq, the corresponding average segment RR intervals are selected. The next step removes from the set of high quality segments any one that has an RRh(k) value close to the corresponding accelerometer segment average interpeak interval (or twice that value in case the ECG sequence reflects only every other major accelerometer peak). (The accelerometer cadence time series may be interpolated to provide values for accelerometer segments that don’t have a high quality sequence and therefore no directly determined cadence value.) The remaining RRh(k) values are passed through a second difference filter to remove outliers.

The next step removes groups of RRh(k) that aren’t sufficiently close in time to enough other surviving RRh(k) values. All RRh(k) that remain are then interpolated so that there is an RR interval curve for all k. For each segment previously eliminated by the above process, the segment’s average RR intervals are examined to determine which is closest to the interpolated RR(k) value. If the closest RR value is within a certain threshold (e.g. 30ms) of the interpolated RR(k) value, it replaces the interpolated /RR(k) value.

### 2.4 Processing Details

All computations were performed on a 2017 Hewlett Packard Laptop with an Intel Core i3-8130U CPU, base frequency 2.20GHz, with 8GB of RAM. The number of candidate peaks (NCP) is set equal to 30. Candidate Peaks must meet broad minimum and maximum peak width criteria.

## 3. Experiment

Four simultaneous chest and waist strap textile ECG recordings were made while a subject walked and jogged on a dirt road for a period of approximately 20-25 minutes after a short period of little or no physical activity just after the initiation of signal acquisition. The dirt road was characterized by uphill, flat, and downhill sections. The four recordings were made on different days. The subject was a 54 year old male. The chest recordings were made with a Hexoskin Smart Shirt while the waist recordings were made with a Zephyr™ Bioharness. Salt water was applied to all textile electrodes before the corresponding garments were worn. The Bioharness was worn so that one textile electrode patch was approximately an inch below and just to the left of the bellybutton, while the other patch was near the front side of the right hip. Athletic tape was layered around the Bioharness to provide a more secure fit. The Bioharness module, worn just to the left of the left textile patch, includes an accelerometer, which produced the accelerometer signals used for processing.

## 4. Results

Figures 5a and 5b show the heart rate of the waist recordings compared to the reference chest recordings. Heart rate was averaged over 5 second segments. The heart rates closely track one another, expect for two approximately 5 minute long recordings in records three and four, shown in Figure 5b. These sections were associated with very poor signal quality, as indicated by the percentage of true beats (reference recording) that occurred in the 30 candidate peaks in the waist recordings. As shown in Figure 5b, during these periods, the percentage of true beats in the candidate set dropped significantly at the outset of these two periods.

**Figure 5a.**
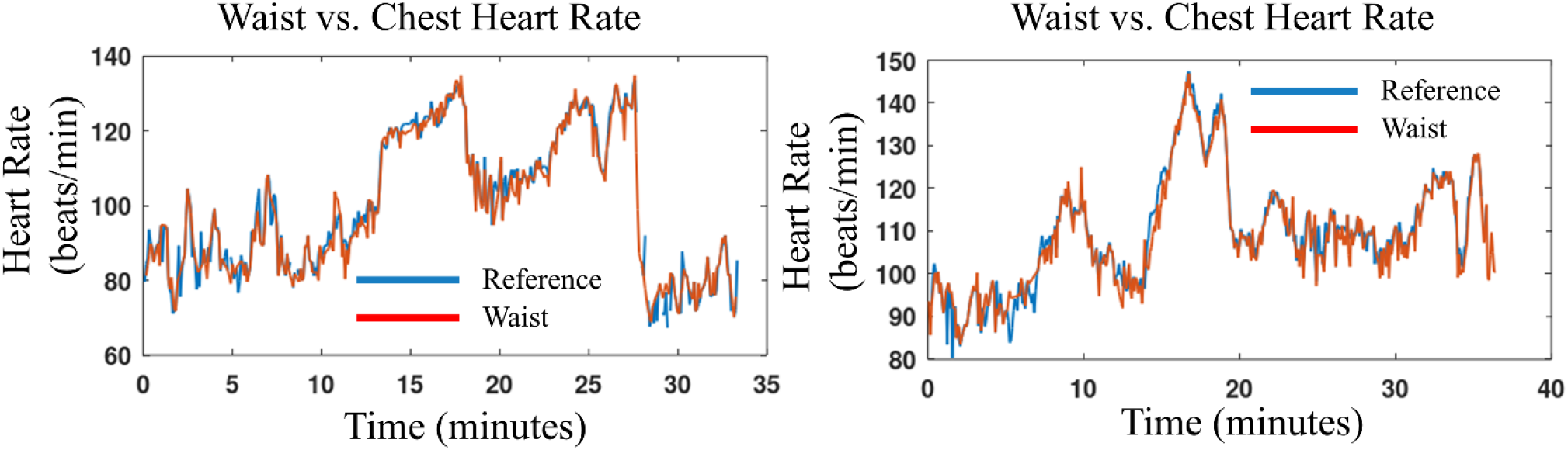
Heart rate while walking and jogging in recordings 1 and 2..

**Figure 5b.**
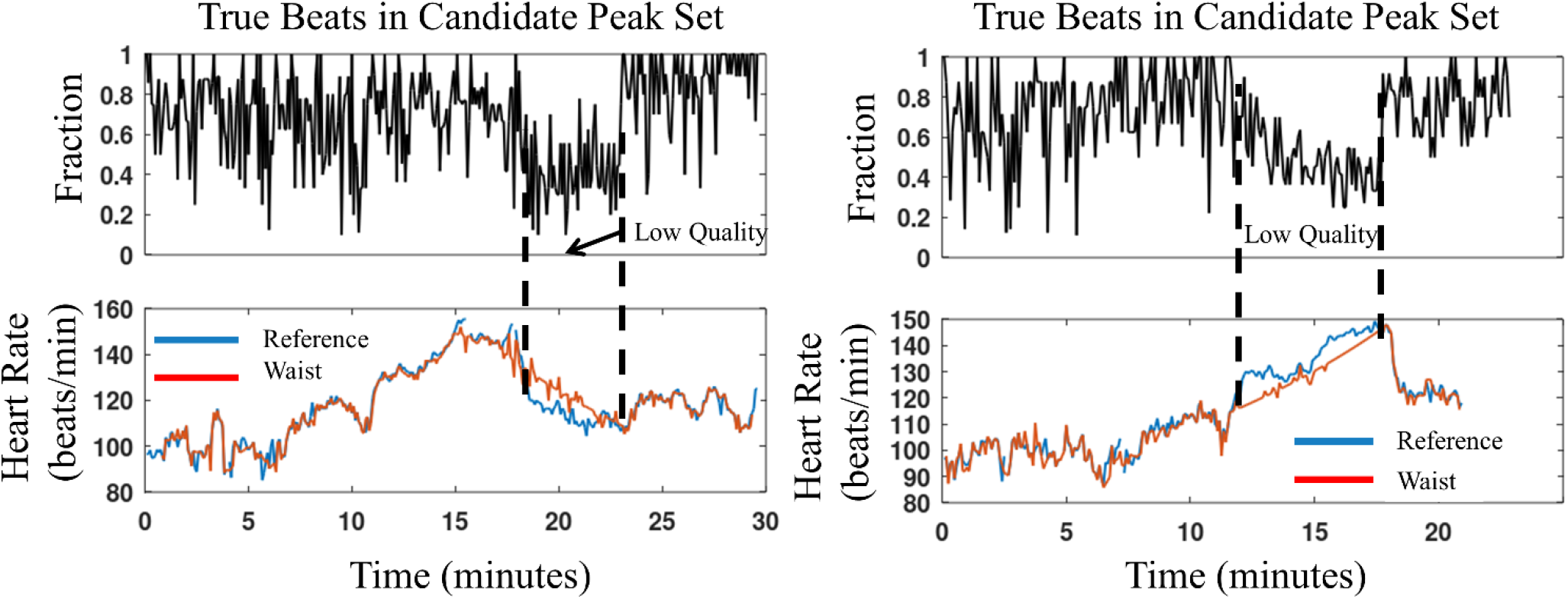
Heart rate while walking and jogging (lower panels) for recordings 3 and 4. The relatively poor performance of the waist recording during an approximately 5 minute period just after jogging in recording 3 (left panels) resulted from the low quality signal, as indicated by the relatively low percentage of true peaks in the set of candidate peaks. A similar signal quality problem occurred while jogging in recording 4.

The first of these two periods occurred just after jogging ceased in recording number 3 (left panels, Figure 5b). Inspection of the signal during this time revealed very high amplitude noise that did not appear to be associated with any motion. This may have been electromyogram noise. The noise was so large that most of the QRS peaks were obliterated, making any sort of detection impossible.

The second these two periods occurred approximately two minutes after jogging started in recording number 4. In this case, the overall signal amplitude did not appear to change much but rather the QRS complexes were frequently indistinct rather than being destroyed by additive noise.

The table below shows the percentage of average heart rate (over 5 second segments) in the waist recordings within 5 and 10 beats/minute of the reference chest recording.

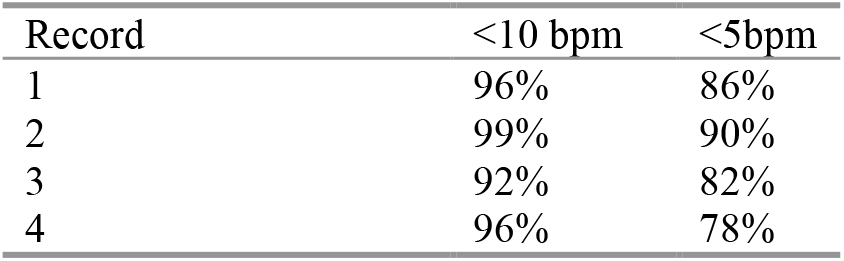

| Record | <10 bpm | <5bpm |
| --- | --- | --- |
| 1 | 96% | 86% |
| 2 | 99% | 90% |
| 3 | 92% | 82% |
| 4 | 96% | 78% |

As described in Section 2.3, the final RR path comprises a combination of high quality segments and segments with RR intervals close to the path defined by interpolating the high quality RR intervals. Together, these two contributing types of segments are referred to “acceptable quality segments.” For acceptable quality segments, the root mean square sum of successive RR interval differences (RMSSD)[8], excluding skipped beats, was calculated for each segment for both the reference and waist recordings. For each segment, the difference between waist and reference RMSSDs was calculated (ΔRMSSD). The table below shows the mean (across segments) ΔRMSSD in each recording.

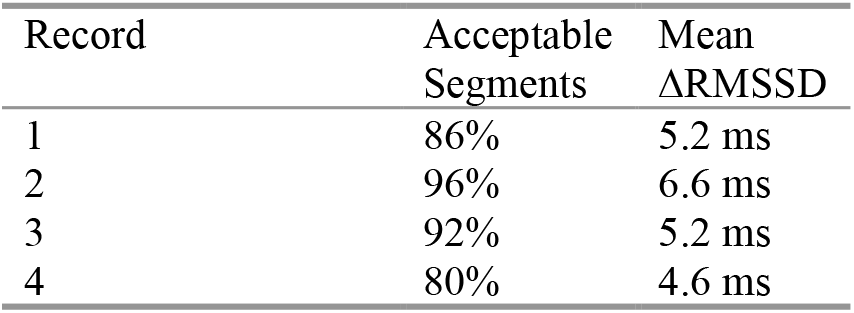

| Record | Acceptable Segments | Mean ΔRMSSD |
| --- | --- | --- |
| 1 | 86% | 5.2 ms |
| 2 | 96% | 6.6 ms |
| 3 | 92% | 5.2 ms |
| 4 | 80% | 4.6 ms |

## 5. Discussion

Figures 5a and 5b show that the waist recordings generally tracked the heart rate of the reference recordings. Given that only one waist ECG channel was available, and that the strap used for the waist recording was not tailored for that purpose, the results point toward the possibility of accurate waist based sinus rhythm detection.

Although motion artifacts were generally most severe during jogging, as expected, the effects of these artifacts were greatly reduced by the peak space cancellation. Conventional cadence measurements tend to be more accurate during jogging compared to walking[9]; in the present case, jogging was typically associated with high quality accelerometer peak sequences and consistent time delays to the corresponding ECG peaks. These artifacts were handled well by the algorithm. However, the motion artifacts need to be kept to the level where they don’t completely obliterate most or all of the QRS complexes, which in turn requires a secure fit of the waistband. Otherwise, motion artifacts will render any sort of QRS detection impossible.

Motion artifact cancellation was not as complete during walking compared to jogging because the accelerometer sequences tended to be relatively lower quality and/or the time delay to the corresponding ECG peaks were not as consistent. Nonetheless, peak cancellation helped with the processing of walking segments. Furthermore, exclusion of RR intervals close to the walking cadence (Section 2.3) was important to the accuracy of the final RR path.

Figure 7 illustrates the somewhat complicated relationship between accelerometer and ECG peaks during walking. Three relatively high amplitude ECG artifact peaks were successfully removed. One true peak was removed as an artifact, as shown by the arrow pointing to the letter B, but this did not prevent detection of the rest of the sequence, including the true peak pointed to by arrow A, which was very close to an appropriately removed accelerometer peak. The ECG peaks pointed to by arrow C were not removed, even though they occurred close in time to an accelerometer peak. A more sophisticated cancellation scheme, perhaps implemented with neural networks, could possibly result in the removal of these peaks without too many unwanted removals of true peaks.

**Figure 7.**
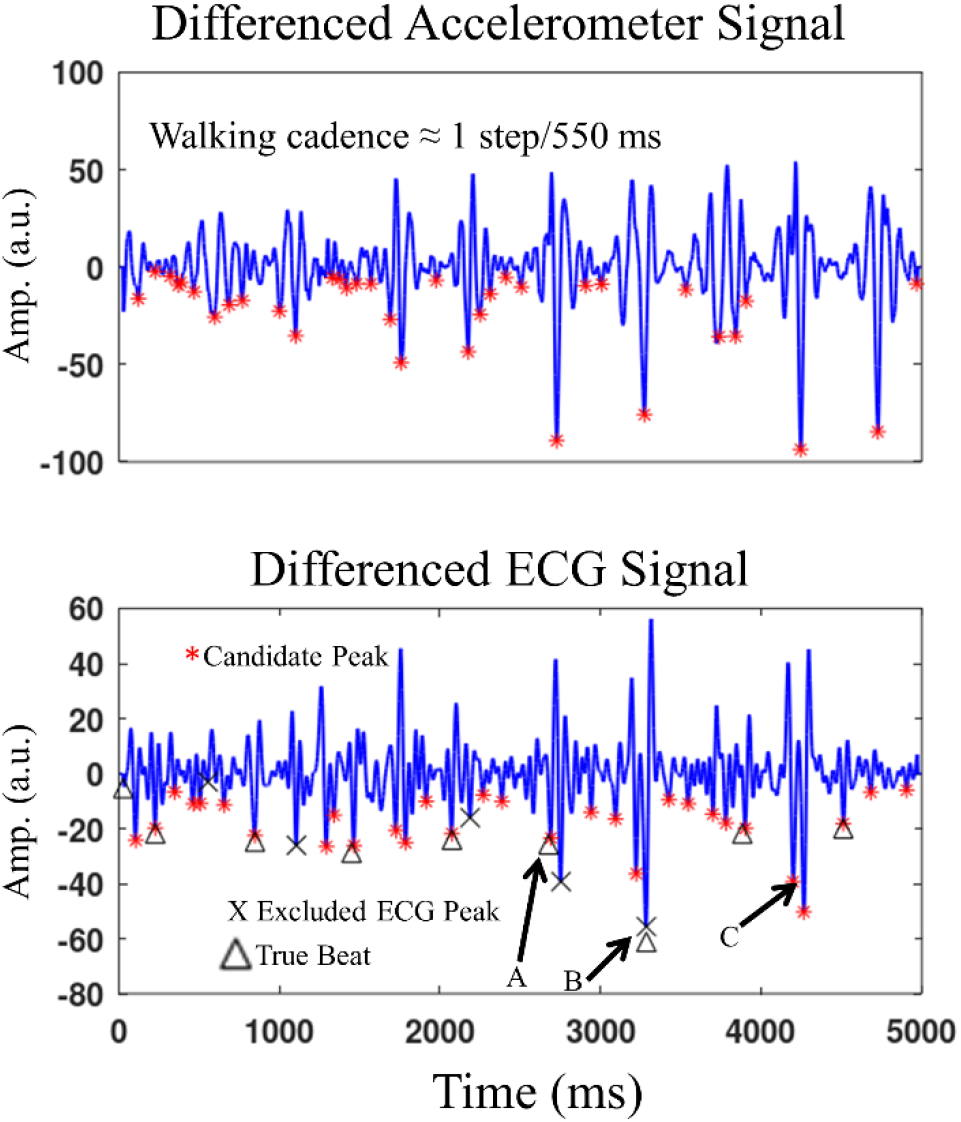
Accelerometer and ECG signals during a 5 second segment recording while the subject was walking at a cadence fairly close to the heart rate. Arrow A points to a correctly detected true peak that is very close to an excluded accelerometer peak. Arrow B points to a true peak that coincided with an excluded accelerometer peak. Arrow C points to artifacts that correlate to an accelerometer peak but that were not marked for exclusion because they did not meet the temporal clustering exclusion criteria.. All true peaks were detected except for the one pointed to by arrow B.

The methodology described herein applies to somewhat regular motions such as walking or jogging. Artifacts that result from more irregular motions (e.g. driving on a rutted dirt road) may require eliminating the temporal regularity score of the accelerometer sequence selection process and instead relying solely on the prominence of the accelerometer peaks. This type of motion will tend to be less of a problem for sinus rhythm detection, but it is still desirable to remove as many artifact peaks as possible, and such elimination may be particularly important for detection of irregular rhythms such as atrial fibrillation[10].

As described above, if a winning ECG sequence (SW) for a particular segment has an RR interval close to that of a corresponding accelerometer sequence, the associated RR interval/segment is not considered high quality for the purpose of determining a final RR segment time series. An alternative approach involves rerunning TEPS on the ECG segment with the peaks corresponding to SW removed.

